## Supplementary figures and tables for "Bacterial exonuclease III expands its enzymatic activities on single-stranded DNA"

**Supplemetary information for**

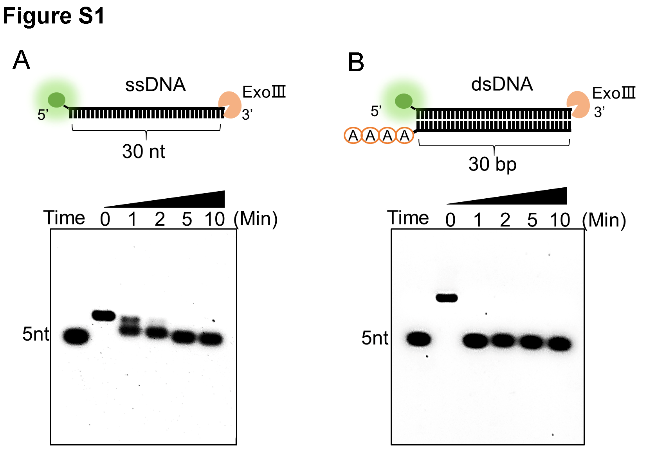

**Supplementary Figure S1.** The digestion process of ExoⅢ on ssDNA and dsDNA substrates was compared. The commercial ExoⅢ (10 U/µl) was incubated with the FAM-labeled ssDNA or dsDNA (10 µM). The reaction products at different timepoints were analyzed by the 6% agarose gel.

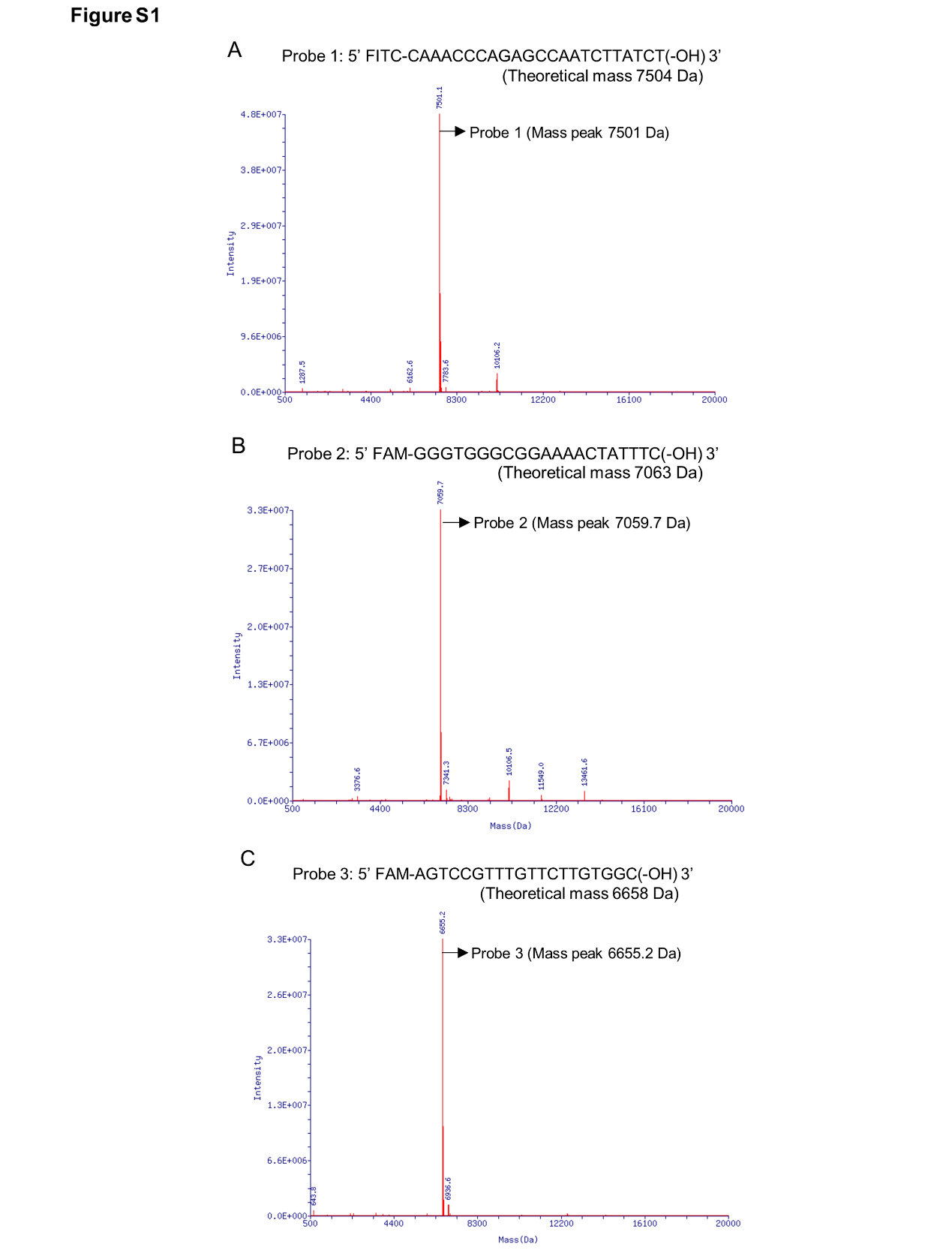

**Supplementary Figure S2.** As controls of the ExoⅢ-treated ssDNAs**,** the mass peaks of ssDNA probes undigested by ExoⅢ were present at **(A)** for Probe 1, **(B)** for Probe 2, and **(C)** for Probe 3. The arrows indicate the major mass peaks produced in Probe 1 (Mass peak = 7501.1 Da, Intensity = 4.79E+07)**,** Probe 2 (Mass peak = 7059.7 Da, Intensity = 3.34E+07), and Probe 3 (Mass peak = 6655.2 Da, Intensity = 3.30E+07). These mass peaks match the theoretical mass of ssDNA Probe 1 (7504 Da), ssDNA Probe 2 (7063.2 Da), and ssDNA Probe 3 (6658 Da). Detailed information on major mass peaks is described in Supplementary information, Table S2.

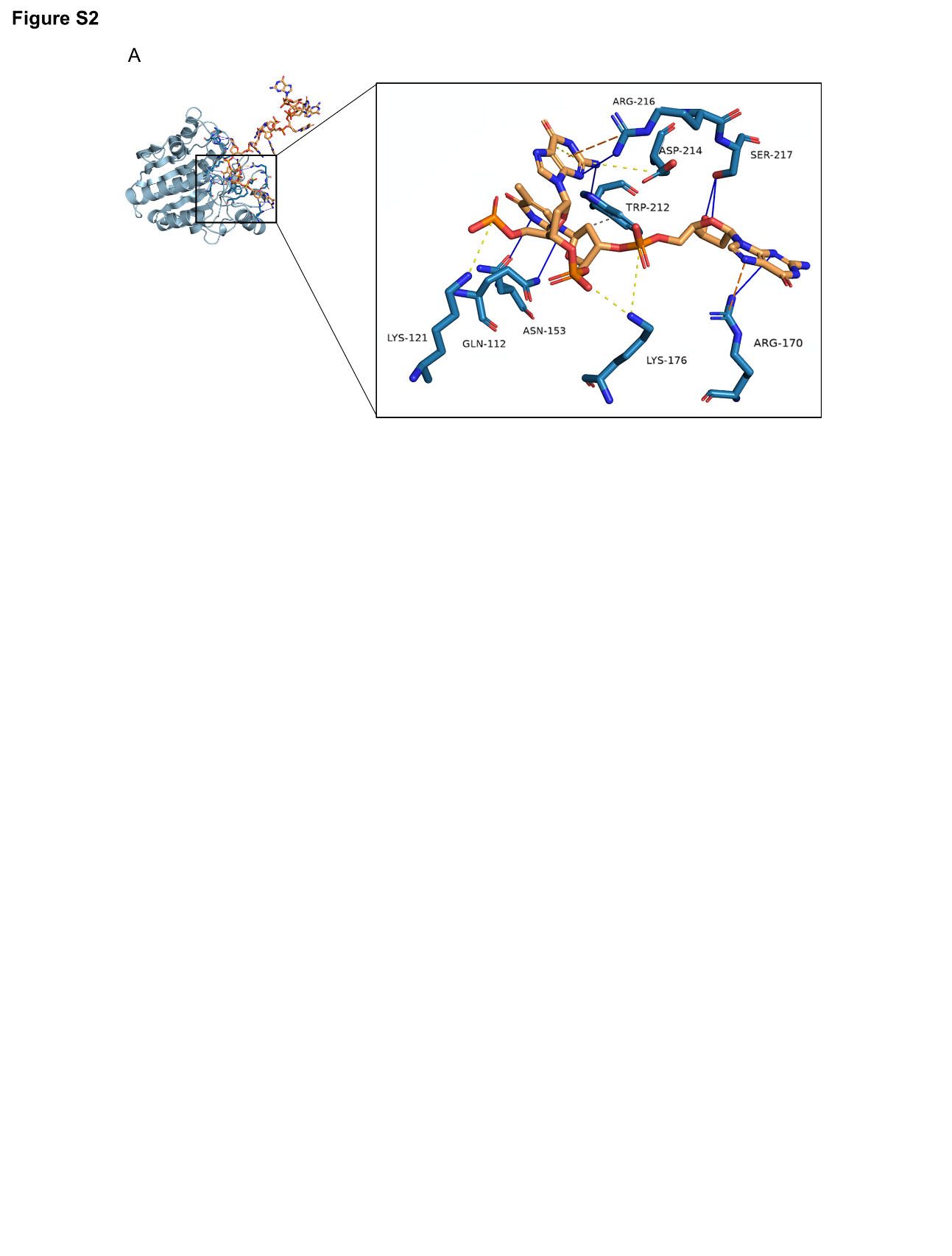

**Supplementary Figure S3.** The molecular docking of ExoⅢ (PDB code: 1AKO) and ssDNA (NDB ID: 1S40) was performed at the HADDOCK docking server (https://wenmr.science.uu.nl/haddock2.4/), and the result was displayed in detail. Residues (SER217, ARG216, ASP214, TRP212, LYS176, ARG170, ASN153, LYS121, and GLN112) were predicted to interact with ssDNA during the enzymatic catalyzation.

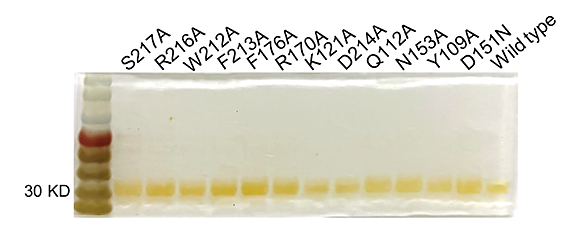

**Supplementary Figure S4.** All purified mutants and wildtype proteins used in the study were examined by the silver-stained SDS-PAGE gel.

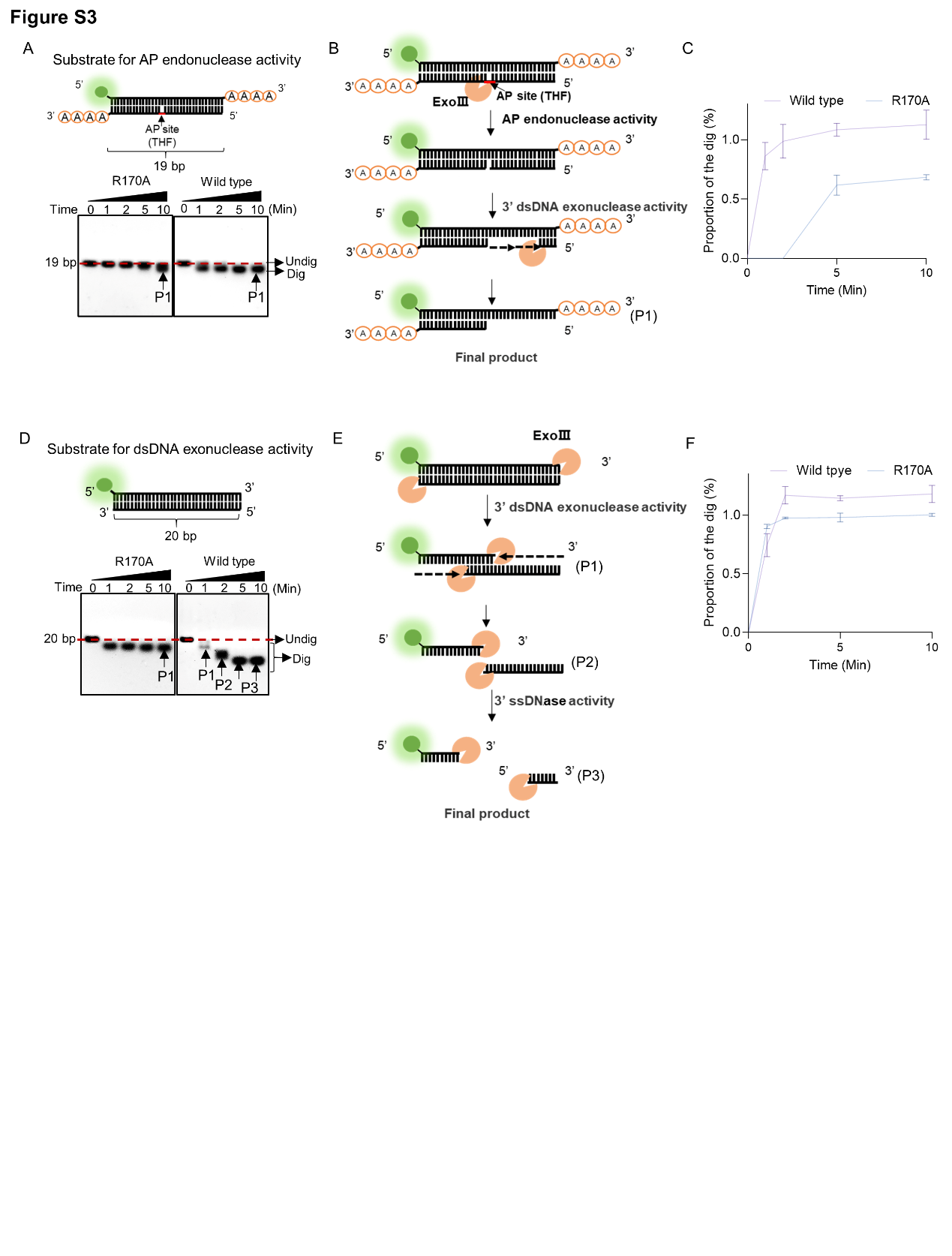

**Supplementary Figure S5.** R170A mutant exhibited a weakened AP-endonuclease activity. **(A)** The dsDNA structure with blunt ends was displayed as a substrate for 3′ exonuclease activity of ExoⅢ. The substrate was digested by R170A and wild type ExoⅢ for 10 min, respectively. Time course analysis of the digestions was presented by gel electrophoresis. **(B)** The digestion process of ExoⅢ on the substrate is diagramed based on the activities identified. **(C)** The Gray intensity of the bands was measured by ImageJ. Then, the average intensity value of three repeats was to calculate the proportion of the digested band by (Intensity of the digested band)/(Intensity of the undigested band at 0 min). **(D)** The dsDNA structure with 3′end A4 was used as the substrate for AP-endonuclease activity. The substrate was incubated with R170A and wild type ExoⅢ for 10 min, respectively. Time course analysis of the digestions was performed. **(E)** The digestion process of ExoⅢ on the substrate is portrayed based on the activities identified. **(F)** The gray intensity of the bands was measured by ImageJ. The average intensity value of three repeats was to calculate the proportion of the digested band by: (Intensity of the digested band)/(Intensity of the undigested band at 0 min).

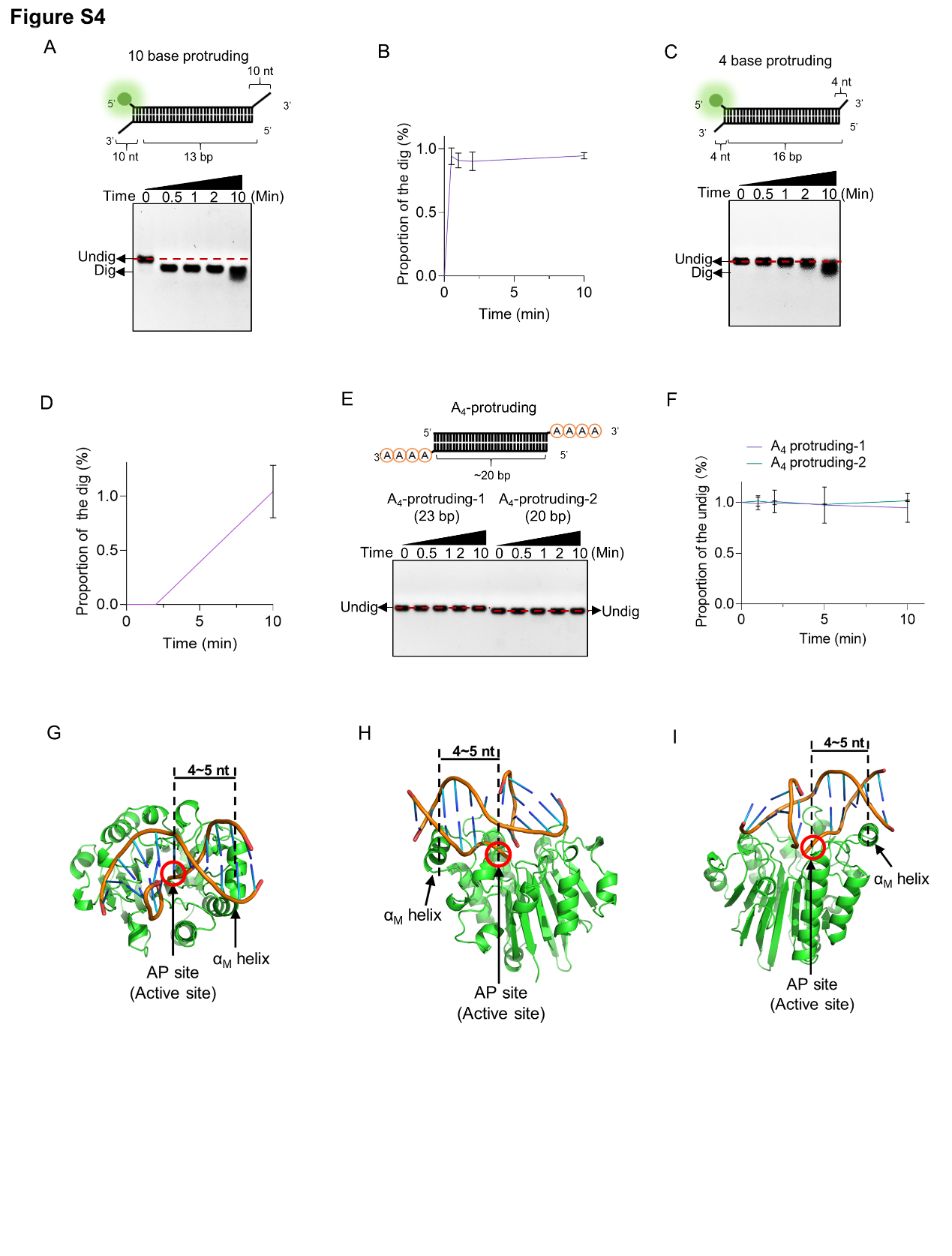
**Supplementary Figure S6.** The dsDNA substrates with 3′ protruding bases (≥ 4 nt) were digested by ExoⅢ. (A) The dsDNA substrate with a 3′ 10-base protruding structure (5 µM) was incubated with ExoⅢ (2.5 µM) for 10 min, and the time course analysis of ExoⅢ digestion was performed. (B) The gray intensity of the bands was determined by ImageJ. The average intensity value of three repeats was to calculate the proportion of the digested band by: (Intensity of the digested band)/(Intensity of the undigested band at 0 min). (C) The dsDNA substrate with a 3′ end 4-base protruding structure was incubated with ExoⅢ for 5 min, and the time course of ExoⅢ digestion was analyzed by gel electrophoresis. (D) The Gray intensity of the bands was determined by ImageJ. The average intensity value of three repeats was to calculate the proportion of the digested band by: (Intensity of the digested band)/(Intensity of the undigested band at 0 min). (E) The dsDNA substrates with 3′ consecutive four-As protruding structure were incubated with ExoⅢ for 10 min, and the time course of ExoⅢ digestion was analyzed by gel electrophoresis. Undig, undigested; dig, digested. (F) The gray intensity of the bands was determined by ImageJ. The average intensity value of three repeats was to calculate the proportion of the undigested band by: (Intensity of the undigested band)/(Intensity of the undigested band at 0 min). The detailed information on these substrates is described in Supplementary Table S4. The αM helix of E. coli is 4~5 bp apart from the active site. The structure of ExoⅢ-dsDNA was obtained in the structure alignment between APE1-dsDNA (PDB ID: 1DE8 and NDB ID: 5WN5) and ExoⅢ (PDB ID: 1AKO) by Pymol. The binding interface between ExoⅢ and dsDNA containing an AP site was present from three view angles: the top (G) and the two sides (H, I).

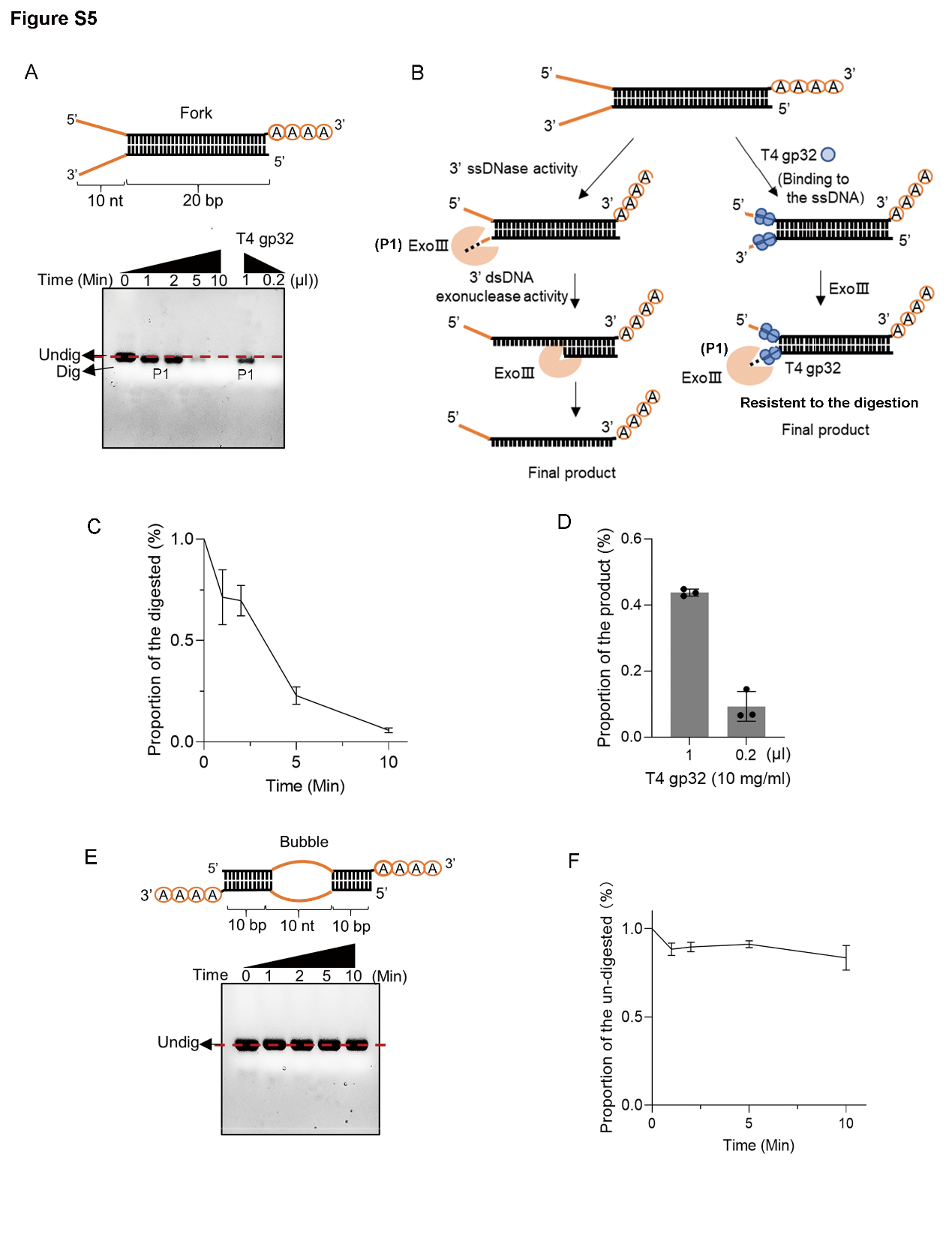

**Supplementary Figure S7.** ExoⅢ digested the fork dsDNA structure with 3' end ssDNA, while the ssDNA binding protein T4 gp32 suppressed the digestion. **(A)** The constitution of fork dsDNA structure is present. The time course analysis of ExoⅢ digestion on the structure was performed, and the products were analyzed by gel electrophoresis. 0.2 or 1 µl ssDNA binding protein T4 gp32 (10 mg/ml) was pre-incubated in the fork structure for 10 min before ExoⅢ digestion. The reaction products were analyzed by gel electrophoresis. **(B)** Based on the activities of ExoⅢ on ssDNA and dsDNA and the protection effect of T4 gp32, the process of enzymatic reaction on the fork dsDNA structure was outlined. **(C)** The digested proportions of ssDNA substrates were calculated by (Gray intensity of the digested band)/(Gray intensity of the undigested band at 0 min) and plotted based on the average value of three repeats. **(D)** The product proportions were calculated by: (Gray intensity of the band with T4 gp32 pre-treatment)/(Gray intensity of the band without T4 gp32 pre-treatment and ExoⅢ digestion at 0 min) and plotted based on the average value of three repeats. **(E)** The constitution of bubble dsDNA structure with 3' end A_4_ is portrayed. The time course analysis of ExoⅢ digestion on the structure was performed, and the products were analyzed by gel electrophoresis. The nucleic acid dye was used in the gel to visualize the reaction products containing the dsDNA duplex. **(F)** The undigested proportions of ssDNA substrates were calculated by (Gray intensity of the undigested band)/(Gray intensity of the undigested band at 0 min) and plotted based on the average value of three repeats.

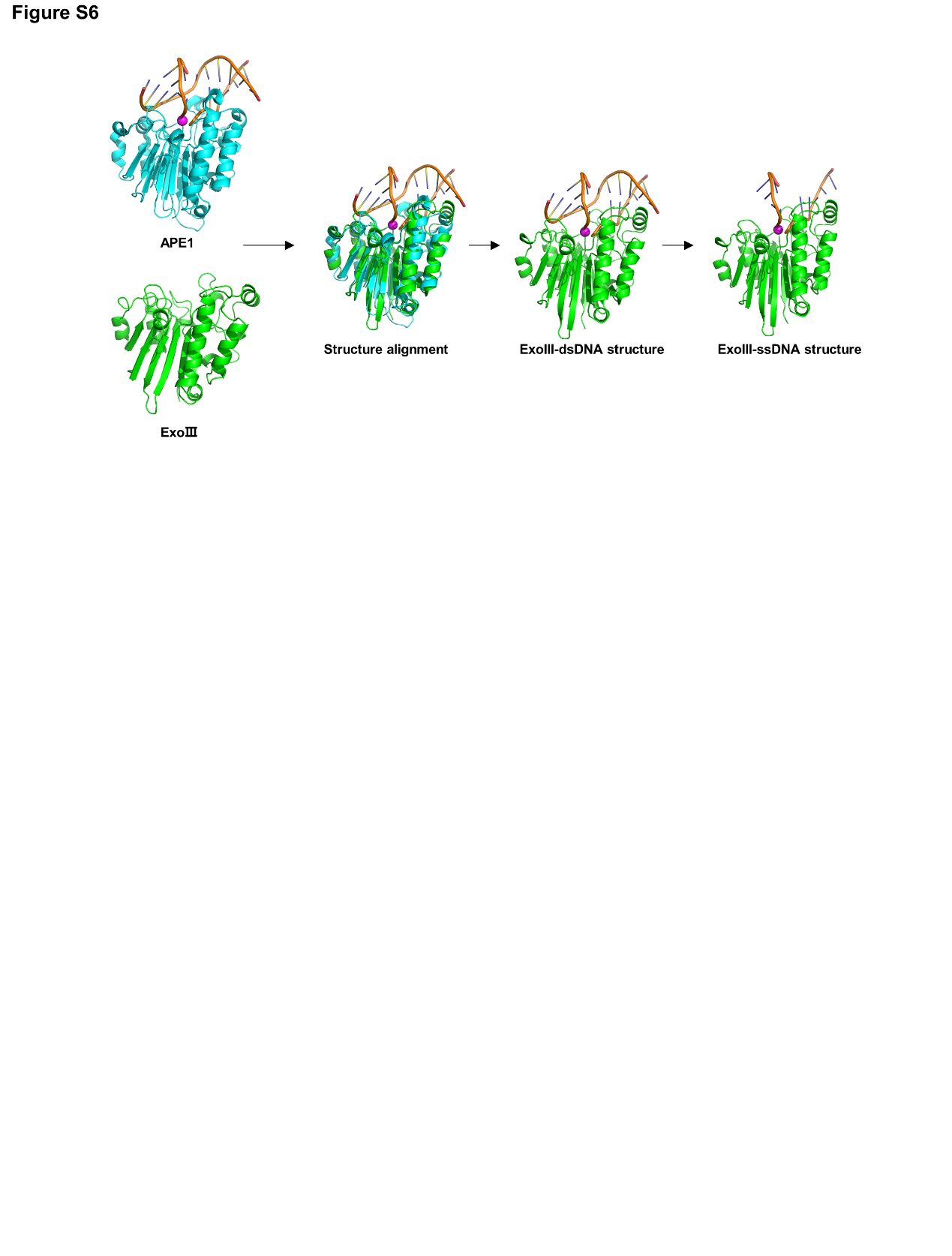

**Supplementary Figure S8.** The residues of ExoⅢ for molecular docking of ExoⅢ and ssDNA were obtained. Structure alignment was performed between ExoⅢ (PDB ID: 1AKO) and APE1-dsDNA (PDB ID: 1DE8 and NDB ID: 5WN5) complex through TM-align (Version 20190822) (https://zhanggroup.org/TM-align/). The two similar homologous structures were perfectly superpositioned to each other. Then, the structure of APE1 and one ssDNA of dsDNA were deleted in the alignment by Pymol, while the other ssDNA that stretches into the active site of ExoⅢ forms the ExoⅢ-ssDNA structure with ExoⅢ. In the structure, residues of ExoⅢ located within 5 Å of ssDNA were identified by Pymol software for molecular docking.

**Supplementary Tables**

| **Supplementary Table S1. The mass spectrometry analysis on the ExoⅢ-treated ssDNA oligos.** | | | | | | | | | |
| --- | --- | --- | --- | --- | --- | --- | --- | --- | --- |
| Sample  (Theoretical mass) | ssDNA without ExoⅢ | | | | ssDNA treated with ExoⅢ | | | | |
|  | Mass peak (Da) | Intensity | Relative (%) | Presumed products (Theoretical mass) | Mass peak (Da) | Intensity | Relative (%) | Presumed products (Theoretical mass) | Structure of the top intensity |
| Probe 1 (7504) | 7501.1 | 4.79E+07 | 100 | FITC-CAAACCCAGAGCCAATCTTATCT (7504) | 2914.9 | 6.31E+06 | 100 | FITC-CAAACCCA (2915.6) | 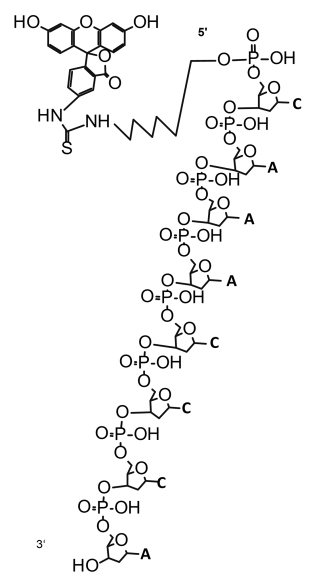 |
|  | 7523.4 | 1.69E+07 | 35.28 | FITC-CAAACCCAGAGCCAATCTTATCT (7504) + Na^+^ | 2601.5 | 4.76E+06 | 75.37 | FITC-CAAACCC (2602.4) |  |
|  | 7546.1 | 1.02E+07 | 21.4 | FITC-CAAACCCAGAGCCAATCTTATCT (7504) + 2Na^+^ | 4047.7 | 1.10E+06 | 17.43 | Undefined |  |
|  | 7567.9 | 3.45E+06 | 7.2 | FITC-CAAACCCAGAGCCAATCTTATCT (7504) + 3Na^+^ | 2312.2 | 1.07E+06 | 17.01 | Undefined |  |
|  | 10106.2 | 3.10E+06 | 6.47 | Undefined | 5395.5 | 6.18E+05 | 9.79 | FITC-CAAACCCAGAGCCAAT (5397.2) |  |
| Probe 2 (7063) | 7059.7 | 3.34E+07 | 100 | FAM-GGGTGGGCGGAAAACTATTTC (7063) | 1287.5 | 5.53E+05 | 100 | Mass peak of ExoⅢ | 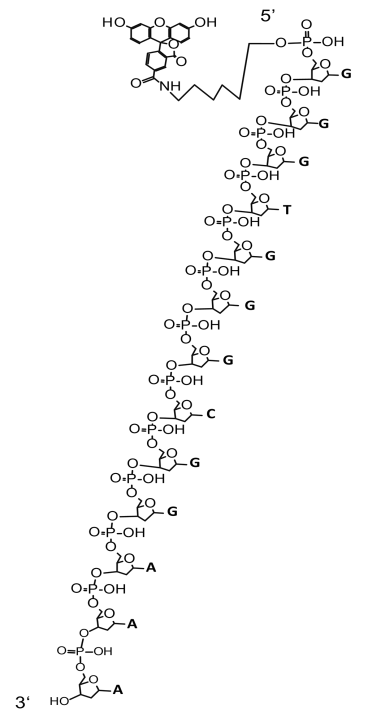 |
|  | 7082.3 | 7.84E+06 | 23.46 | FAM-GGGTGGGCGGAAAACTATTTC (7063) + Na+ | 627.8 | 3.35E+05 | 60.59 | Mass peak of ExoⅢ or TT (P) (626.4) |  |
|  | 7104.6 | 2.95E+06 | 8.84 | FAM-GGGTGGGCGGAAAACTATTTC (7063) + 2Na^+^ | 13646.1 | 2.01E+05 | 36.26 | Undefined |  |
|  | 10106.5 | 2.27E+06 | 6.8 | Undefined | 10107.2 | 1.97E+05 | 35.57 | Undefined |  |
|  | 7341.3 | 1.20E+06 | 3.58 | Undefined | 1979.4 | 1.92E+05 | 34.81 | Undefined |  |
|  | 13461.6 | 1.07E+006 | 7.94 | Undefined | 4639.6 | 1.59E+05 | 28.74 | FAM-GGGTGGGCGGAAA (4641.6) |  |
|  | 11549.0 | 6.24E+005 | 6.55 | Undefined | 6655.3 | 1.55E+05 | 28.11 | Undefined |  |
|  | 3376.6 | 4.57E+005 | 2.88 | Undefined | 1291.6 | 1.45E+05 | 26.29 | Mass peak of ExoⅢ or GCGG/GGCG/GGGC (P) (1293.8) |  |
|  | 7481.5 | 3.82E+005 | 5.67 | Undefined | 9126.5 | 1.02E+05 | 18.42 | Undefined |  |
|  |  |  |  |  | 957.7 | 1.02E+05 | 18.39 | AAA (P) (957.6) |  |
| Probe 3 (6658) | 6655.2 | 3.30E+07 | 100 | FAM-AGTCCGTTTGTTCTTGTGGC (6658) | 2936.9 | 8.36E+06 | 100 | FAM-AGTCCGTT (2937.6) | 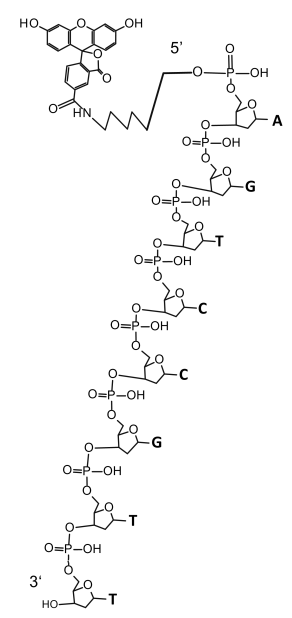 |
|  | 6677.5 | 1.06E+07 | 32.18 | FAM-AGTCCGTTTGTTCTTGTGGC (6658) + Na^+^ | 3240.9 | 2.77E+06 | 33.1 | FAM-AGTCCGTTT (3241.8) |  |
|  | 6699.9 | 6.00E+06 | 18.15 | FAM-AGTCCGTTTGTTCTTGTGGC (6658) + 2Na^+^ | 2632.7 | 2.34E+06 | 27.99 | FAM-AGTCCGT (2633.4) |  |
|  | 6721.6 | 1.85E+06 | 5.59 | Undefined | 2328.7 | 8.11E+05 | 9.69 | FAM-AGTCCG (2329.2) |  |
|  | 6936.6 | 1.29E+06 | 3.91 | Undefined | 643.8 | 4.59E+05 | 5.48 | Undefined |  |
| ExoⅢ |  |  |  |  | 1287.6 | 5.44E+05 | 100 | Undefined |  |
|  |  |  |  |  | 627.8 | 3.45E+05 | 63.38 | Undefined |  |
|  |  |  |  |  | 2639.4 | 1.91E+05 | 35.09 | Undefined |  |
|  |  |  |  |  | 1291.6 | 1.47E+05 | 27.04 | Undefined |  |
|  |  |  |  |  | 19174.8 | 1.36E+05 | 25.08 | Undefined |  |
| Note that P indicates the oligo with a phosphate group. A (dAMP) = 331.2 g/mol, T (dTMP) = 322.2 g/mol, G (dGMP) = 347.2 g/mol, and C (dCMP) = 307.2 g/mol. Molecular weight of the 5′ end FITC part (C27H25N2O5S) of the reaction product is 489 g/mol, and the 5′ end FAM part (C_27_H_24_NO_6_) is 458 g/mol. The red indicates the zone of three identical bases where the digestion of ExoⅢ stalled. | | | | | | | | | |

| **Supplementary Table S2. The** **embedded use of ssDNA in various ExoⅢ-associated diagnostic platforms** | | | | | |
| --- | --- | --- | --- | --- | --- |
| Classification by the ExoⅢ activity used in the methods | Diagnostic platforms | Targets | Functions of ExoⅢ | Involved nucleic acid aptamer | Published |
| dsDNA exonuclease-based detection | ExoⅢ-aided electrochemiluminescencence techniques | miRNA-21 (a cancer biomarker) | Recognize and degrade dsDNA to ssDNA for triggering subsequent reaction | The ssDNA oligos and hairpin dsDNA with 3′ end protruding ssDNA or dsDNA with the blunt end | 2023 ^44^ |
|  | ExoⅢ combined with quantum dots. | HIV and HBV virus |  |  | 2021 ^45^ |
|  | ExoⅢ combined with nanoparticles | p53 gene |  |  | 2021 ^46^ |
|  | ExoIII integrated with the microfluidic platform. | Manganese superoxide dismutase gene |  |  | 2020 ^47^ |
|  | ExoⅢ facilitated chemiluminescence techniques. | Synthetic DNA target |  |  | 2014^48^ |
| AP-endonuclease based detections | ExoⅢ integrated with Recombinase Polymerase Amplification (RPA) | Aeromonas salmonicida | Recognize and cleave the AP site on dsDNA for releasing fluorescence signal. | Primers and a ssDNA oligo containing AP site labeled with fluorescence and quencher groups | 2022^35^ |
|  |  | Burkholderia cepacia |  |  | 2023^49^ |
|  |  | Elizabethkingia miricola |  |  | 2022^50^ |
|  |  | Porcine parvovirus |  |  | 2017^51^ |
|  |  | Dengue virus |  |  | 2015^52^ |
|  |  | SARS-CoV-2 |  |  | 2020^53^ |

| **Supplementary Table S3. An updated list for ExoⅢ activities** | | | | |
| --- | --- | --- | --- | --- |
| Substrates | Structures | Activities | Products | Key references |
| dsDNA | With 3′ phosphonate | 3′ phosphatase | Phosphate | ^10^ |
|  | Containing AP | Endonuclease | ssDNA gap on DNA | ^7^ |
|  | Natural DNA | Exonuclease | Phosphate, 5′ mononucleotides, ssDNA | ^11^ |
| RNA | RNA/DNA hybrid | RNase H | Mono-ribonucleotide, ssDNA | ^54^ |
| ssDNA | Containing AP | Active (less than dsDNA) | Oligos | ^55^ |
|  | Nonstructural ssDNA | Exonuclease and endonuclease activities | Mononucleotide, Oligos | This study |

| **Supplementary Table S4. A list of used sequences in the study.** | | | | |
| --- | --- | --- | --- | --- |
| Oligos | Sequences | 5′ Modifications | 3′ Modifications | Used |
| FQ reporter | TTATT | FAM (labeled at phosphonate) | BHQ1 (labeled at phosphonate) | Fig. 1, 4, 5, 6 |
| T_1_-labeled reporter | T_1_TATT | FAM (at T_1_ base) | BHQ1 (at phosphonate) | Fig. 1 |
| Base-labeled FQ reporter | TTATT | FAM (at T_1_ base) | BHQ1 (at T base) | Fig. 1 |
| Probe 1 | CAAACCCAGAGCCAATCTTATCT | FITC (at phosphonate) | None (-OH) | Fig. 2, 3, 4 |
| Probe 2 | GGGTGGGCGGAAAACTATTTC | FAM (at phosphonate) | None (-OH) | Fig. 2, 3, 4 |
| Probe 3 | AGTCCGTTTGTTCTTGTGGC | FAM (at phosphonate) | None (-OH) | Fig. 2, 3, 4, 5, 6 |
| Activator-S for Cas12a trans-cleavage activity | TTTCAACAGCACATGCAGAATCAT | None (-OH) | None (-OH) | Fig. 1, 2 |
| Activator-AS for Cas12a trans-cleavage activity | ATGATTCTGCATGTGCTGTTGAAA | None (-OH) | None (-OH) | Fig. 1, 2 |
| crRNA | GGUAAUUUCUACUAAGUGUAGAUAACAGCACAUGCAGAAUCAU | None (-OH) | None (-OH) | Fig. 1, 2 |
| A_20_ | AAAAAAAAAAAAAAAAAAAA | FAM (at phosphonate) | None (-OH) | Fig. 4 |
| C_20_ | CCCCCCCCCCCCCCCCCCCC | FAM (at phosphonate) | None (-OH) | Fig. 4 |
| T_20_ | TTTTTTTTTTTTTTTTTTTT | FAM (at phosphonate) | None (-OH) | Fig. 4 |
| Substrate-S for endonuclease activity | AAGATTGGCTCTGGGTTTGAAAA | FAM (at phosphonate) | None (-OH) | Supplementary Fig. S3 |
| Substrate-AS for endonuclease activity | CAAACCCAG[THF]GCCAATCTTAAAA | None (-OH) | None (-OH) | Supplementary Fig. S3 |
| Substrate-S for exonuclease activity | CAAACCCAGAGCCAATCTTATCT | FAM (at phosphonate) | None (-OH) | Supplementary Fig. S3 |
| Substrate-AS for exonuclease activity | AGATAAGATTGGCTCTGGGTTTG | None (-OH) | None (-OH) | Supplementary Fig. S3 |
| 10 base protruding structure-S | CAAACCCAGAGCCAATCTTATCT | FAM (at phosphonate) | None (-OH) | Supplementary Fig. S4 |
| 10 base protruding structure-AS | GGCTCTGGGTTTGATCGATCGATCG | None (-OH) | None (-OH) | Supplementary Fig. S4 |
| 4 base protruding structure | CAAACCCAGAGCCAATCTTATCT | FAM (at phosphonate) | None (-OH) | Supplementary Fig. S4 |
| 4 base protruding structure-AS | AAGATTGGCTCTGGGTTTGATCG | None (-OH) | None (-OH) | Supplementary Fig. S4 |
| A_4_ protruding structure-1-S | CAAACCCAGAGCCAATCTTAAAA | None (-OH) | None (-OH) | Supplementary Fig. S4 |
| A_4_ protruding structure-1-AS | AAGATTGGCTCTGGGTTTGAAAA | None (-OH) | None (-OH) | Supplementary Fig. S4 |
| A_4_ protruding structure-2-S | AGTCCGTTTGTTCTTGAAAA | None (-OH) | None (-OH) | Supplementary Fig. S4 |
| A_4_ protruding structure-2-AS | CAAGAACAAACGGACTAAAA | None (-OH) | None (-OH) | Supplementary Fig. S5 |
| Bubble-S | TGTGAGTTTTGAGCGTGGCGTGCTGGAGCAAAAA | None (-OH) | None (-OH) | Supplementary Fig. S5 |
| Bubble-AS | TGCTCCAGCAATCGATCGATAAAACTCACAAAAA | None (-OH) | None (-OH) | Supplementary Fig. S5 |
| Fork-S | TGTGAGTTTTGAGCGTGGCGTGCTGGAGCAAAAA | None (-OH) | None (-OH) | Supplementary Fig. S5 |
| Fork-AS | TGCTCCAGCACGCCACGCTCATCGATCGAC | None (-OH) | None (-OH) | Supplementary Fig. S5 |
| 3′flap on dsDNA-S-1 | AGTCCGTTTGTTCTTGTGGC | FAM (at phosphonate) | None (-OH) | Fig. 8 |
| 3′flap on dsDNA-S-2 | TAAGATTGGCAAAAA | None (-OH) | None (-OH) | Fig. 8 |
| 3′flap on dsDNA-AS | GCCAATCTTACAAACGGACTAAAAA | None (-OH) | None (-OH) | Fig. 8 |
| Note that S stands for sense strand, and AS stands for antisense strand. | | | | |
